# The Increasing Authorship trend in Neuroscience: A scientometric analysis across 11 countries

**DOI:** 10.1101/2024.01.24.577031

**Authors:** Ann Paul, Mariella Segreti, Pierpaolo Pani, Emiliano Brunamonti, Aldo Genovesio

**Affiliations:** Department of Physiology and Pharmacology, Sapienza University, Rome, Italy; Behavioral Neuroscience PhD Program, Sapienza University, Rome, Italy

**Keywords:** Neuroscience, Authorship, G10 countries

## Abstract

Previous studies have demonstrated an increasing trend of the number of authors across various fields over the years. This trend has been attributed to the necessity for larger collaborations and, at times, to ethical issues regarding authorship attribution. Our study focuses on the evolution of authorship trends in the field of Neuroscience. We conducted our analysis based on a dataset containing 576,647 neuroscience publications produced from 2000 to 2022, focusing on the publications within the Group of ten (G10) countries. Using a matrix-based methodology, we extracted and analyzed the average number of authors per country. Our findings reveal a consistent rise in authorship across all G10 countries over the past two decades. Italy emerged with the highest average number of authors, while the Netherlands stood out for experiencing the most significant increase, particularly in the last decade. The countries with the lowest number of authors per publication were the USA, UK, and Canada.

## 1. Introduction

Collaboration is essential in academic research, promoting the exchange of ideas and the integration of diverse expertise and innovative methodologies. Today’s authors also embrace “altruism” in research, recognizing that collective efforts pave the path to more entries on their publication roster (Wuchty et al., 2007).

Since the late 1980s, numerous European and non-European countries have implemented national systems to monitor, assess, and evaluate the research performance of their scientific workforce (Hicks, 2012; Baccini et al., 2019, Baccini & Petrovich, 2023). A prominent feature of these research evaluation systems is the emphasis on quantitative indicators, recognized as pivotal science policy tools (Ingwersen & Larsen, 2014). As a result, in recent years, various scientometric indicators, centered on the number of publications, citations, or their combination (such as the h-index), have been adopted in academic evaluation systems. The seminal work by Baccini et al. (2019) shed light on this phenomenon, offering insights into the escalation of the citations in all fields especially focusing on the G10 countries with special concern for the Italian authors. Also, Fox and colleagues (2016) revealed a significant correlation between the two variables, investigating data collected from 32 ecology journals spanning the years 2009 to 2012. Additionally, papers with a greater number of authors are more prone to potential self-citations, as noted by Larivière et al. (2014), and benefit from a larger network of colleagues who may cite the paper (Borsuk et al., 2009; Fox et al., 2016).

While extensive research has focused on citation metrics, the evolution in time of the number of authors, as a quantitative indicator, has been overlooked compared to other indicators. Over the years, in fact, a limited number of studies (Sampson, 1995;Abt, 2007; Aboukhalil, 2014); Camargo Jr. & Coeli, 2012; Gu et al., 2017; An et al., 2020) have explored the number of authors in articles in specific fields or journals. These studies revealed a significant paradigm shift in research dynamics within the scientific community. This shift is characterized by an increase in collaborative efforts, exemplified by the growing number of authors per paper. For instance, in the field of Physics, Zuckerman and Merton’s investigation into the archives of The Physical Review between 1948 and 1956 revealed a prevalence of single-authored papers, constituting over half of the 14,512 manuscripts examined (Zuckerman & Merton, 1971; Sampson, 1995). However, by the 1980s, the landscape had shifted, with articles featuring three or more authors surpassing the 50% mark (Sampson, 1995). In astronomy a similar pattern was observed, with early 20th-century papers predominantly single-authored, contrasting sharply with the 4.9 ± 0.7 average authors per paper in 2005. In 15 other science fields, the average of the number of authors in 2005 ranged from 3.0 in mathematics to 7.4 in cardiology, with an overall average of 5.5 ± 0.4 authors per paper (Abt, 2007). However, Abt’s work indicates that while single-authored papers are decreasing, they are not likely to vanish any time soon. In his opinion, the reduction of single-authored papers is indicative of the changes in the nature of scientific inquiry and will not affect all types of research. Aboukhalil (2015) has described the rising trend in authorship from 1913 to 2014. According to Aboukhalil, the number of authors for publication has increased over 5-fold over the last century and is projected to reach an average of 8 authors per paper by 2034. Finally, Fortunato et al. (2018) have shown that scientific teams are growing in size, with an average increase of 17% per decade.

Within the influential domain of the Group of Ten (G10) countries, the landscape of publication is undergoing a discernible metamorphosis (Baccini et al., 2019) This transformation determines a surge in the number of authors per paper, capturing the attention of the scholarly community. What was once the domain of individual expertise and singular contributions has evolved into a collective endeavor, drawing together multiple researchers, and experts across disciplines and geographical boundaries. These findings revealed a growing trend in the number of authors, but systematic analyses of this phenomenon are limited. Previous studies (Sampson, 1995; Abt, 2007; Gu et al., 2017; An et al., 2020) have explored specific sectors or journals without a comparison across countries. In this study we chose to study the temporal trend of the number of authors for the first time in neuroscience across 11 countries (G-10 countries + 1) through a systematic examination of 272 neuroscience journals and the analysis of 576,647 publications.

## 2. Methods

The dataset for the analysis consisted of the entire 2000-2022 neuroscience publications from the Web of science (WoS, www.webofscience.com). A total of 272 Neuroscience journals included in this analysis (see Supplementary information) were indexed in the source website. Out of 12,80,000 publications (including articles, editorial materials, reviews, and letters) that were downloaded, 5,76,647 publications were published in the G-10 countries, that are: Belgium, Canada, France, Germany, Italy, Japan, the Netherlands, Sweden, Switzerland, United Kingdom, and United States. Following the criteria adopted by Baccini et al. (2019), these countries were selected as they contributed to 61.2% of the world output in terms of scientific publications in the years 2000-2016 (Baccini et al., 2019).

### 2.1. Data Analysis

After having computed the total number of authors for publication, along with the respective country of the corresponding author and the year of publication, we averaged the number of authors per year. To measure changes in authorship patterns with time, we used the delta measure by calculating the differences between two sub-periods (2000-2011 & 2012-2022), for a quantitative assessment of the variations. We conducted a two-way ANOVA (Country X Sub-period) for testing if the number of authors significantly differed between countries and periods of time. Tukey-Kramer post-hoc tests were used for further investigations. The statistical analysis was performed in MATLAB R2021 (www.mathworks.com).

## 3. Results

Figure 1 illustrates the temporal trend of authorship in the field of Neuroscience for the eleven target countries from 2000 to 2022. Despite variations in individual countries, a consistent overall pattern emerged, with a regular growth in authorship in all countries examined. We computed the average number of authors across all journals, spanning the period from 2000 to 2022. As illustrated in Table 1, we found that Italy holds the lead in terms of authorship, having the highest number of authors. In contrast, Canada ranked the lowest near the United States and UK.

**Table 1.** Average number of authors with standard deviation across all the journals, spanning the period from 2000 to 2022 and in the two sub-periods: 2000-2011 and 2012-2022 are reported for each country. The delta value between the two sub-periods is also reported.

| Country | Publications | Average No. of authors (2000-2022) | Average No. of authors (2000-2011) | Average No. of authors (2012-2022) | Δ Delta |
| --- | --- | --- | --- | --- | --- |
| Italy | 39120 | 6.55 (0.92) | 5.89 (0.42) | 7.27 (0.76) | 1.38 |
| Japan | 46400 | 6.10 (0.57) | 5.66 (0.34) | 6.59 (0.29) | 0.93 |
| France | 28872 | 5.82 (1.00) | 5.02 (0.36) | 6.69 (0.68) | 1.67 |
| Germany | 54778 | 5.58 (0.83) | 4.95 (0.37) | 6.26 (0.62) | 1.31 |
| Netherlands | 19572 | 5.55 (1.06) | 4.74 (0.35) | 6.43 (0.84) | 1.69 |
| Belgium | 8561 | 5.47 (0.72) | 4.90 (0.25) | 6.08 (0.53) | 1.18 |
| Sweden | 10418 | 5.26 (0.80) | 4.63 (0.33) | 5.96 (0.51) | 1.33 |
| Switzerland | 11642 | 5.25 (0.71) | 4.73 (0.35) | 5.81 (0.84) | 1.08 |
| United Kingdom | 53102 | 4.84 (0.89) | 4.15 (0.33) | 5.59 (0.64) | 1.45 |
| United States | 264479 | 4.82 (0.84) | 4.17 (0.34) | 5.53 (0.59) | 1.37 |
| Canada | 39703 | 4.55 (0.84) | 3.90 (0.28) | 5.27 (0.62) | 1.37 |
| *Mean G10 Countries* | 576647 | 5.44 (0.59) | 4.79 (0.61) | 6.14 (0.59) | 1.34 (0.23) |

**Figure 1.**
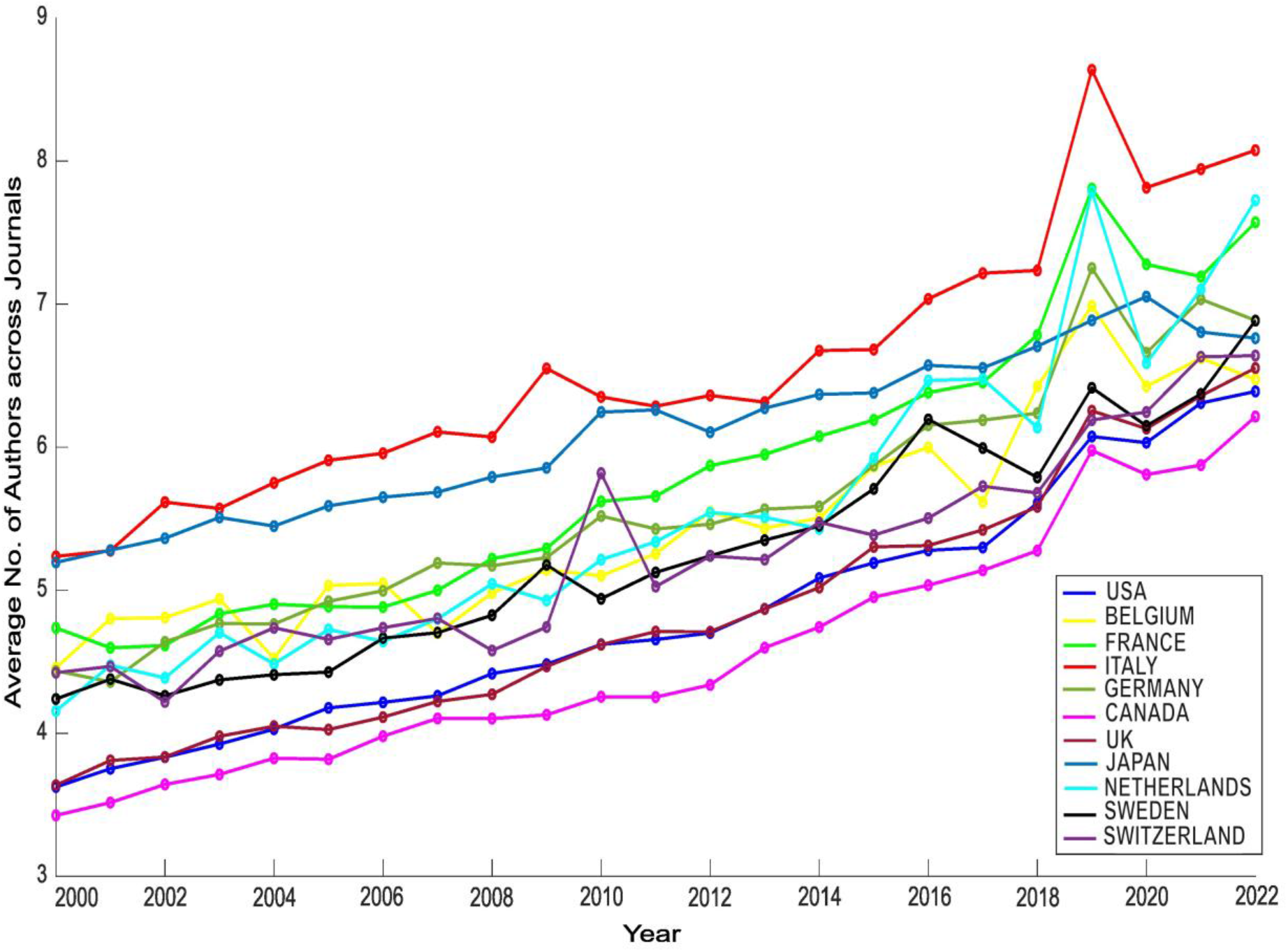
Temporal trend of authorship in the field of Neuroscience for the eleven target countries from 2000 to 2022.

For estimating whether the average number of authors differed between countries and its variation in time, we conducted a within-country two-way ANOVA with country and sub-periods as factors. We found that each factor independently exerted a significant effect, as evidenced by the statistical significance of both the country factor (F (10,242) = 31, p < 0.0001) and the sub-period factor (F (1,242) = 403, p < 0.0001). The interaction between the two factors was not statistically significant (F (10,263) = 1, p = 0.4). Tukey-Kramer post-hoc comparisons performed identified significant differences within the eleven countries. We found that the publications coming from Italy had, in both sub-periods, a significantly higher number of authors than all other countries (all p < 0.001 except for Japan). In addition, the significant effect of the sup-period factor indicates that over all countries the number of authors per publication increased from the decade 2000-2011 to 2012-2022. To assess the consistency of growth in the number of authors in each country, we computed the variations (deltas) in authorship for each country between the two sub-periods: 2000-2011 and 2012-2022. We followed the same methodology used for the calculation of inwardness by Baccini and colleagues (2019). We split the two periods across 2010 because that was the year when the reform of the university was introduced in Italy, which changed the criteria for evaluating researchers’ productivity. Table 1 shows the variations (deltas) in authorship for each country. Across countries we observed an average increase of 1.34 (1.23) authors from the decade 2000-2011 to 2012-2022. Examining single countries, we observed that the Netherlands stood out with the largest authorship delta (1.69), establishing itself as the country with the highest authorship delta and showing the most significant growth in the number of authors between the first and second periods.

Figure 1 shows a pronounced spike in 2019, which is hard to explain considering only the increasing trend of the number of authors. To understand it we tested whether that spike could be accounted for by an increase of publications with an exceptionally high number of authors in 2019. Examining the publications with more than 40 authors, we observed a distinct pattern of authorship in 2019 with publications with up to 400 authors. Some of these publications reported among the authors not only authors with standard authorship, but also included clinics and laboratories, attributing authorship to all affiliated members.

Considering the 169 publications produced in 2019, we found that a high proportion of those with more than 40 authors focused on specific research fields such as, ageing brain studies mostly related to neurodegenerative diseases (23%) or Genomic projects (18%) that might require a larger than average scale of collaboration for the acquisition or elaboration of data. This is the reason why, during the retrieval of publications from Web of Science, some included up to 400 authors.

We performed a new analysis that excluded all the publications exceeding this 40 authors threshold, by removing 806 (0.14% of the total) publications from the total number of publications of all the countries in the years examined. Figure S1 shows the new graph that lacks the 2019 spike.

The same ANOVA was performed including only the publications below 40 authors. As with the full dataset of publications we found that each factor independently exerted a significant effect, as evidenced by the statistical significance of both the ‘Country’ factor (F (10,220) =50, p < 0.0001) and the sub-period factor (F (1,220) = 514, p < 0.0001) with no significant interaction between the two (F (10,241) = 1, p = 0.5). Even with this restricted selection, Italian publications had a significantly higher number of authors than all other countries in both the sub-periods, (Tukey-Kramer post-hoc, p <0.05).

## 4. Discussion

Extensive research has explored indicators such as citation metrics to evaluate the researcher’s production (Hicks, 2012; Baccini et al., 2019). The studies on the trend of the number of authors have focused on specific sectors and not considered all the journals in those fields (Zuckerman & Merton, 1971; Sampson, 1995; Abt, 2007). Specific fields have been taken into consideration in previous studies by assessing the number of authors per research area rather than per country. For instance, the study by Fernandes and Monteiro (2017) divided Computer Science into 17 constituent areas to analyze the number of authors per area and witnessed in all areas an increase in the average number of authors per paper across all decades (1954–2014). An and colleagues (2020) adopted a comparable approach, examining clinical journals across different medical specialties reporting the highest increase in the number of authors in articles compared to review articles and case reports. These studies collectively highlighted the pervasive trend in the increase of the number of authors and the evolving nature of authorship practices across various scientific disciplines.

While previous studies focused on specific sectors within disciplines, a systematic examination across a specific discipline, considering all journals and conducting a cross-country analysis is still lacking. Our research addressed this gap, conducting a systematic analysis of 1,280,000 neuroscience publications to analyze the trend of the number of authors within the entire neuroscience sector. Furthermore, our study evaluated the similarity of growth in authorship across various temporal periods and targeted countries, exploring whether distinct national patterns emerged.

Our analysis revealed that, among the G10 countries under consideration for the period 2000 to 2022, Italy boasts the highest average number of authors in the field of neuroscience. Additionally, the Netherlands showed the greatest increase in authorship over this timeframe. The reasons behind Italy having the highest number of authors and the Netherlands experiencing the most significant increase in the second subperiod (2012-2022) are aspects that merit further exploration.

Analyzing the type of authorship that characterized the 2019 spike we found that it was marked by a higher than usual number of authors which could include not only standard authors but also the entire clinic or laboratory, attributing authorship to all affiliated members. We found that the primary area of interest of these publications was on specific topics such as neurodegenerative disorders and genetics. In the context of neurodegenerative disorders like Alzheimer’s disease and multiple sclerosis, most publications were based on longitudinal studies performed with the help of collaborations with clinics and hospitals (Schilling et al., 2019). Considering genomics, the explanation for the high number of authors has to be found in the nature of the research that involves large-scale projects with numerous collaborators (Brazel et al., 2019) given the complexity of genomic studies. Why these studies had a pick of publication in 2019 requires further investigation.

In addressing the heterogeneity of authorship, we propose a classification into three categories of authorship based on the nature of authorship. The first category, referred to as the standard authors, includes the authors that are listed in the publication and in the repository where the publication is stored, presenting a comprehensive list with their names (e.g., Koopmans et al., 2019).

The second category includes collaborators. Repositories and archives such as PubMed and Europe PMC, first lists the standard authors and then includes a section called “Collaborators,” which should be expanded to read the contributors who, while not directly engaged in the writing process, have played a non-specified role in the publications (e.g., Barger et al., 2019).

The third category involves consortiums; we refer to this third category of authors as consortium authors. In certain publications and repositories, not only individual authors are listed, but also entire consortiums are introduced without specifying the identity of each contributor (e.g., van der Lee et al., 2019). Despite the absence of an explicit listing of the individual contributors within the publication or repository, Web of Science adopts a comprehensive approach, by recognizing all members of the consortium as authors. This practice of recognizing authorship in this way can result in publications being credited to as many as hundreds of authors. How the second and third categories of authorship should be considered in the evaluation process of a researcher’s productivity should become a topic of discussion to prevent evaluation algorithms from making decisions that we might not be aware of.

Our findings raise several questions. First of all, we have explored the wide-ranging field of neuroscience without dividing it by subfields. In computer science Fernandez and Montero (2017) showed that the number of authors varies across subfields, with emerging fields such as bioinformatics and multimedia having more authors. We anticipate that similar differences may exist in neuroscience, warranting upcoming analysis to explore this aspect.

Future studies should also extend this analysis beyond the G10 countries as done by Baccini and Petrovich (2023) in the study of self-citations. Indeed, Baccini et al. (2023) demonstrated that Italy is the sole G10 country experiencing a recent surge in self-citations diverging from the other countries since 2010. The year 2010 marks the initiation of a government reform (Law 240/2010) in the Italian university system when algorithms were introduced to reward researchers based on the number of citations they received. While it is challenging considering only these countries to understand why Italy is the only country with this increasing trend, an expanded analysis of 50 countries revealed that all the other countries (Colombia, Egypt, Indonesia, Iran, Italy, Malaysia, Pakistan, Romania, Russian Federation, Saudi Arabia, Thailand, and Ukraine) with a trend as Italy have implemented analogous science policies, rewarding individual citations in the evaluation of researcher performance.

Thus, a more extensive comparative approach would contribute to a better understanding of the global dynamics of authorship growth, not only in neuroscience but also in other fields. Future research should also explore more in-depth regional differences within countries, such as variations between the north, center, and south of Italy, as well as distinctions between public and private universities or institutes.

Future studies are needed to understand the dynamics underlying the differences in the number of authors among countries. Firstly, does the variation in the number of authors indicate distinct criteria for inclusion in a publication, considering the level of commitment and time contributed by each author? Secondly, does this difference between countries depend on the inclusion or not of specific categories of authors as undergraduate students or the heads of departments or institutes (Eisenberg et al., 2014)?

For example, a deeper analysis has been provided by Gu and colleagues (2017) on the trends in the specialty of hand surgery. They found a shift in the academic qualifications of first authors over this period. There was a marked increase in the proportion of first authors holding an MD/PhD, PhD, master’s, or bachelor’s degree since 1985 whereas there was a decrease in the proportion of first authors solely holding an MD during the same timeframe. They also reported an increase in manuscripts authored by women over the past 30 years. These findings underscore the importance of replicating similar in-depth analyses in other fields, such as neuroscience.

An essential consideration revolves around whether all listed authors in a publication share equal responsibility for the publication or if the burden falls primarily on the first author when facing ethical questions from the review board. Examining the three classes of authorship, as described by Rennie et al. (1994), the ‘grafters’—individuals listed at the end of the authorship roll, who contribute minimally to the project—may exhibit a lack of commitment. In instances of suspicion or ethical concerns, these names, as highlighted by Solomon (2009) and Bennett et al. (2003), might be prone to fading away and evading accountability. A clear example of this phenomenon is represented by the case of Dr. John Darsee, who was exposed for fabricating medical data (Solomon, 2009). When Darsee’s practices were discovered, his former co-authors did not want to take responsibility even if they accepted to be listed as co-authors (Broad, 1983). This vanishing of the co-authors, who were reluctant to share equal responsibility for the published work, suggests minimal or no contribution to the study (Solomon, 2009). Several studies in multiple fields have reported the presence of honorary authors, as in the Darsee case, defined as individuals who did not contribute or only contributed marginally to the work, highlighting differences between countries (Eisenberg et al., 2014; Marušić et al., 2011) or based on the level of the journal and the specific health field (Aliukonis et al., 2020). In a survey conducted in the biomedical field, it was found that 33.4% of the participating authors reported adding authors to their manuscripts who did not contribute significantly to the work (Al-Herz et al., 2014). When asked why authors who did not contribute were added, in 39.4% of cases, sadly, the reason fell under the category of friendship, returning a favor, or assisting a colleague in obtaining a promotion. Therefore, the number of authors on a publication transcends a simple quantitative measure and, in a larger than expected proportion of cases, indicates an ethical violation.

We should ask whether an increase in the number of authors can be considered as the response of some researchers to an evaluation system that is increasingly reliant on numerical indicators, as proposed by Baccini et al. (2023), which does not correct for the number of authors.

Typically, the competition for academic positions (“publish or perish”) adopts criteria such as the number of publications without taking into account, at least in countries like Italy the number of authors. To get competitive funding for research, a robust publication record featuring numerous publications possibly in high impact factor journals is a prerequisite (Van Arensbergen et al., 2014; Waaijer, 2017). More importantly, the criteria used to evaluate candidates for research positions and career advancements provide an advantage for researchers engaged in more collaborative efforts (Waaijer, 2017). The intentional increase in the number of authors might not only aim to accumulate more publications but also strategically boost citations through their inclusion (Wuchty et al., 2007; Sin, 2011).

In the Italian system, not only the number of publications but also the number of citations together with the h-index are measured to evaluate the researchers’ productivity. The competition for advancement in academia in Italy is, in fact, based on the achievement of a national scientific qualification (ASN), that measure, following the criteria established by an Italian government agency (ANVUR), the scientific productivity in the researcher specific disciplinary sector (SSD) counting number of authors, citations, and using the h-index. Abramo & D’Angelo (2015) reported a positive correlation between the number of authors and the number of citations, specifically in the Italian publications in the neuroscience category during the period 2004-2010 similar to Fox et al. (2016). Considering all the factors that impact collaboration, it emerges that the number of authors is, at least in part, associated with utilitarian considerations that should be considered when defining the criteria for evaluation of the careers of the researchers. What can be done to address this issue? Let us consider the h-index which is a measure of the impact of a scientist’s publications, developed by Hirsch (2005) and it serves as one of the metrics used to evaluate a scientist’s production. This index is calculated as the highest number of papers by a researcher that have been cited h or more times. Since its inception, Hirsch himself made it clear that this index required some form of normalization to account for the number of authors. Despite various proposed corrections of the index over the years (Schreiber, 2008; Galam, 2011), none have been implemented in evaluating researchers’ careers, at least in Italy. The implementation of corrections can rely on very simple algorithms as shown in the correction of the citations, such as the author contribution score (Zerem, 2017), which calculates by assigning the total number of points to the first author, half of the total number of points to the corresponding author (if not the first author), and equally distributing the remaining half points among the other authors. The decision to ignore this issue in the valuation system is puzzling and raises questions in itself. We conclude by questioning how we can contrast this ethical issue when, who is at the head of the institutions and who should offer solutions benefits the most from honorary authorship, especially in the European countries as highlighted by the study of Eisenberg et al. (2014).

## Supporting information

Figure S1 and list of the journals

