## Supplementary material for "The Increasing Authorship trend in Neuroscience: A scientometric analysis across 11 countries": Figure S1 and list of the journals

**SUPPLEMENTARY INFORMATION**

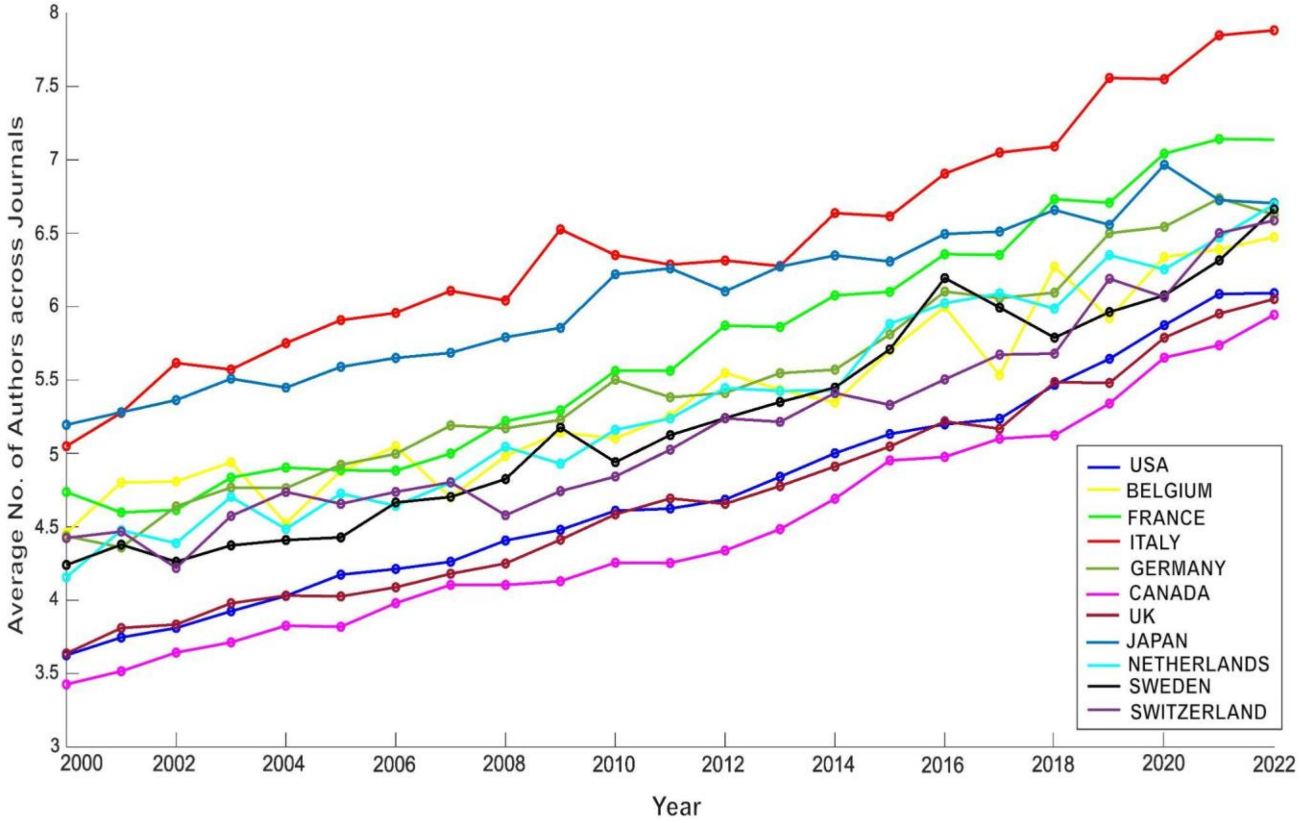


**Figure S1.** Temporal trend of authorship in the field of Neuroscience for the eleven target countries from 2000 to 2022 with publications below 40 authors.

**LIST OF THE JOURNALS**

**Selected Category: NEUROSCIENCES.**
**Year: 2022.**
**Selected Category Schema: WOS.**
**Total number: 272.**
**Copyright (c) 2024 Clarivate.**

- Acs chemical neuroscience
- Acta neurobiologiae experimentalis
- Acta neurologica belgica
- Acta neuropathologica
- Acta neuropathologica communications
- Acta neuropsychiatrica
- Actas espanolas de psiquiatria
- Acupuncture & electro-therapeutics research
- Alzheimers research & therapy
- Annals of clinical and translational neurology
- Annals of neurology
- Annual review of neuroscience
- Annual review of vision science
- Archives italiennes de biologie
- Arquivos de neuro-psiquiatria
- Asn neuro
- Audiology and neuro-otology
- Autonomic neuroscience-basic & clinical
- Behavioral and brain functions
- Behavioral and brain sciences
- Behavioral neuroscience
- Behavioural brain research
- Behavioural pharmacology
- Biological cybernetics
- Biological psychiatry
- Biological psychiatry-cognitive neuroscience and neuroimaging
- Bipolar disorders
- Bmc neuroscience
- Brain
- Brain and behavior
- Brain and cognition
- Brain and language

- Brain behavior and evolution
- Brain behavior and immunity
- Brain connectivity
- Brain impairment
- Brain injury
- Brain pathology
- Brain research
- Brain research bulletin
- Brain sciences
- Brain stimulation
- Brain structure & function
- Brain topography
- Cellular and molecular neurobiology
- Cephalalgia
- Cerebellum
- Cerebral cortex
- Ceska a slovenska neurologie a neurochirurgie
- Chemical senses
- Chemosensory perception
- Clinical autonomic research
- Clinical eeg and neuroscience
- Clinical neurophysiology
- Clinical psychopharmacology and neuroscience
- Cns & neurological disorders-drug targets
- Cns neuroscience & therapeutics
- Cognitive affective & behavioral neuroscience
- Cognitive computation
- Cognitive neurodynamics
- Cognitive neuroscience
- Cognitive systems research
- Cortex
- Current alzheimer research
- Current neurology and neuroscience reports
- Current neuropharmacology
- Current neurovascular research
- Current opinion in behavioral sciences
- Current opinion in neurobiology
- Current opinion in neurology
- Developmental cognitive neuroscience
- Developmental neurobiology
- Developmental neuroscience
- Dialogues in clinical neuroscience
- Encephale-revue de psychiatrie clinique biologique et therapeutique

- Eneuro
- Epilepsia open
- European journal of neurology
- European journal of neuroscience
- European journal of pain
- European neurology
- European neuropsychopharmacology
- Experimental brain research
- Experimental neurobiology
- Experimental neurology
- Fluids and barriers of the cns
- Folia neuropathologica
- Frontiers in aging neuroscience
- Frontiers in behavioral neuroscience
- Frontiers in cellular neuroscience
- Frontiers in computational neuroscience
- Frontiers in human neuroscience
- Frontiers in integrative neuroscience
- Frontiers in molecular neuroscience
- Frontiers in neural circuits
- Frontiers in neuroanatomy
- Frontiers in neuroendocrinology
- Frontiers in neuroinformatics
- Frontiers in neurology
- Frontiers in neurorobotics
- Frontiers in neuroscience
- Frontiers in synaptic neuroscience
- Frontiers in systems neuroscience
- Gait & posture
- Genes brain and behavior
- Glia
- Hearing research
- Hippocampus
- Human brain mapping
- Human movement science
- Ideggyogyaszati szemle-clinical neuroscience
- Ieee transactions on cognitive and developmental systems
- International journal of developmental neuroscience
- International journal of neuropsychopharmacology
- International journal of neuroscience
- International journal of psychophysiology
- Jaro-journal of the association for research in otolaryngology
- Journal of alzheimers disease

- Journal of cerebral blood flow and metabolism
- Journal of chemical neuroanatomy
- Journal of clinical neurophysiology
- Journal of clinical neuroscience
- Journal of cognitive neuroscience
- Journal of comparative neurology
- Journal of comparative physiology a-neuroethology sensory neural and behavioral physiology
- Journal of computational neuroscience
- Journal of electromyography and kinesiology
- Journal of headache and pain
- Journal of integrative neuroscience
- Journal of mathematical neuroscience
- Journal of molecular neuroscience
- Journal of motor behavior
- Journal of musculoskeletal & neuronal interactions
- Journal of neural engineering
- Journal of neural transmission
- Journal of neurochemistry
- Journal of neurodevelopmental disorders
- Journal of neuroendocrinology
- Journal of neuroengineering and rehabilitation
- Journal of neurogenetics
- Journal of neuroimmune pharmacology
- Journal of neuroimmunology
- Journal of neuroinflammation
- Journal of neurolinguistics
- Journal of neuromuscular diseases
- Journal of neuropathology and experimental neurology
- Journal of neurophysiology
- Journal of neuropsychiatry and clinical neurosciences
- Journal of neuroscience
- Journal of neuroscience methods
- Journal of neuroscience research
- Journal of neurotrauma
- Journal of neurovirology
- Journal of pain
- Journal of parkinsons disease
- Journal of physiology-london
- Journal of pineal research
- Journal of psychiatry & neuroscience
- Journal of psychopharmacology
- Journal of psychophysiology

- Journal of sleep research
- Journal of stroke & cerebrovascular diseases
- Journal of the history of the neurosciences
- Journal of the international neuropsychological society
- Journal of the neurological sciences
- Journal of the peripheral nervous system
- Journal of vestibular research-equilibrium & orientation
- Learning & memory
- Metabolic brain disease
- Molecular and cellular neuroscience
- Molecular autism
- Molecular brain
- Molecular neurobiology
- Molecular neurodegeneration
- Molecular pain
- Molecular psychiatry
- Motor control
- Multiple sclerosis journal
- Muscle & nerve
- Nature and science of sleep
- Nature human behaviour
- Nature neuroscience
- Nature reviews neuroscience
- Network neuroscience
- Network-computation in neural systems
- Neural computation
- Neural development
- Neural networks
- Neural plasticity
- Neural regeneration research
- Neurobiology of aging
- Neurobiology of disease
- Neurobiology of learning and memory
- Neurobiology of stress
- Neurochemical journal
- Neurochemical research
- Neurochemistry international
- Neurocirugia
- Neurodegenerative diseases
- Neuroendocrinology
- Neuroendocrinology letters
- Neurogastroenterology and motility
- Neuroimage

- Neuroimaging clinics of north america
- Neuroimmunomodulation
- Neuroinformatics
- Neurologic clinics
- Neurological research
- Neurological sciences
- Neurological sciences and neurophysiology
- Neurology india
- Neurology-neuroimmunology & neuroinflammation
- Neuromolecular medicine
- Neuromuscular disorders
- Neuron
- Neuropathology
- Neuropathology and applied neurobiology
- Neuropeptides
- Neuropharmacology
- Neurophotonics
- Neurophysiologie clinique-clinical neurophysiology
- Neurophysiology
- Neuropsychobiology
- Neuropsychologia
- Neuropsychological rehabilitation
- Neuropsychology
- Neuropsychology review
- Neuropsychopharmacology
- Neuroreport
- Neuroscience
- Neuroscience and biobehavioral reviews
- Neuroscience bulletin
- Neuroscience letters
- Neuroscience research
- Neuroscientist
- Neurotherapeutics
- Neurotoxicity research
- Neurotoxicology
- Neurotoxicology and teratology
- Npj parkinsons disease
- Npj science of learning
- Nutritional neuroscience
- Pain
- Pharmacology biochemistry and behavior
- Progress in neuro-psychopharmacology & biological psychiatry
- Progress in neurobiology

- Psychiatric genetics
- Psychiatry and clinical neurosciences
- Psychoneuroendocrinology
- Psychopharmacology
- Psychophysiology
- Purinergic signalling
- Restorative neurology and neuroscience
- Reviews in the neurosciences
- Seizure-european journal of epilepsy
- Sleep
- Sleep and biological rhythms
- Sleep medicine reviews
- Social cognitive and affective neuroscience
- Social neuroscience
- Somatosensory and motor research
- Stereotactic and functional neurosurgery
- Stress-the international journal on the biology of stress
- Synapse
- Translational neurodegeneration
- Translational neuroscience
- Translational stroke research
- Trends in cognitive sciences
- Trends in neurosciences
- Vision research
- Visual neuroscience
- Zhurnal vysshei nervnoi deyatelnosti imeni i p pavlova
